## Supplementary Table 1 for "Quantifying dynamic facial expressions under naturalistic conditions"

**Supplementary Information**

| **Short description** | **Detailed description** | **Start time (seconds)** |
| --- | --- | --- |
| Happiness | Cute dog saying “I love you”  Young girl on singing show | 0 |
| Surprise | Angry office worker breaks telephone  Man surprised by loud bang | 40 |
| Disgust | Man eats beetle larva | 70 |
| Fear | Large snake swallows a pig | 96 |
| Sadness | Starvation, funerals | 125 |
| Fear | Crocodile show ends with crocodile snapping on man’s hand | 199 |

**Supplementary Table 1.** Videos shown to participants in the DISFA dataset.

| Action unit | Description | Associated emotion |
| --- | --- | --- |
| **1** | Inner Brow Raiser | Sadness, surprise, fear |
| **4** | Brow Lowerer | Sadness, fear, anger |
| **6** | Cheek Raiser | Happiness |
| **7** | Lid Tightener | Fear, anger |
| **9** | Nose Wrinkler | Disgust |
| **10** | Upper Lip Raiser |  |
| **12** | Lip Corner Puller | Happiness |
| **14** | Dimpler |  |
| **15** | Lip Corner Depressor | Sadness, disgust |
| **17** | Chin Raiser | Disgust |
| **20** | Lip Stretcher | Fear |
| **23** | Lip Tightener | Anger |
| **25** | Lips Part |  |
| **26** | Jaw Drop | Surprise, fear |

**Supplementary Table 2.** Facial action units used in our study, with corresponding time series extracted with OpenFace(1). The third column shows emotions conventionally associated with each action unit, in the Emotional Facial Action Coding System(2).

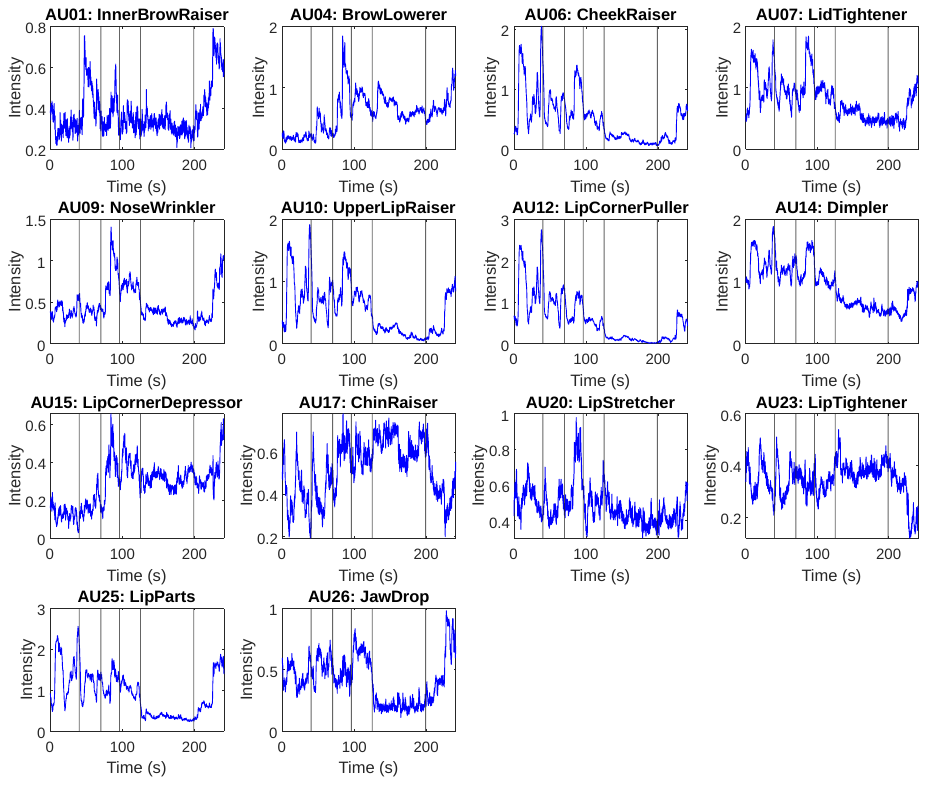

**Supplementary Figure 1.** Mean of raw time series, DISFA dataset. Vertical lines demarcate video clips described in Supplementary Table 1.

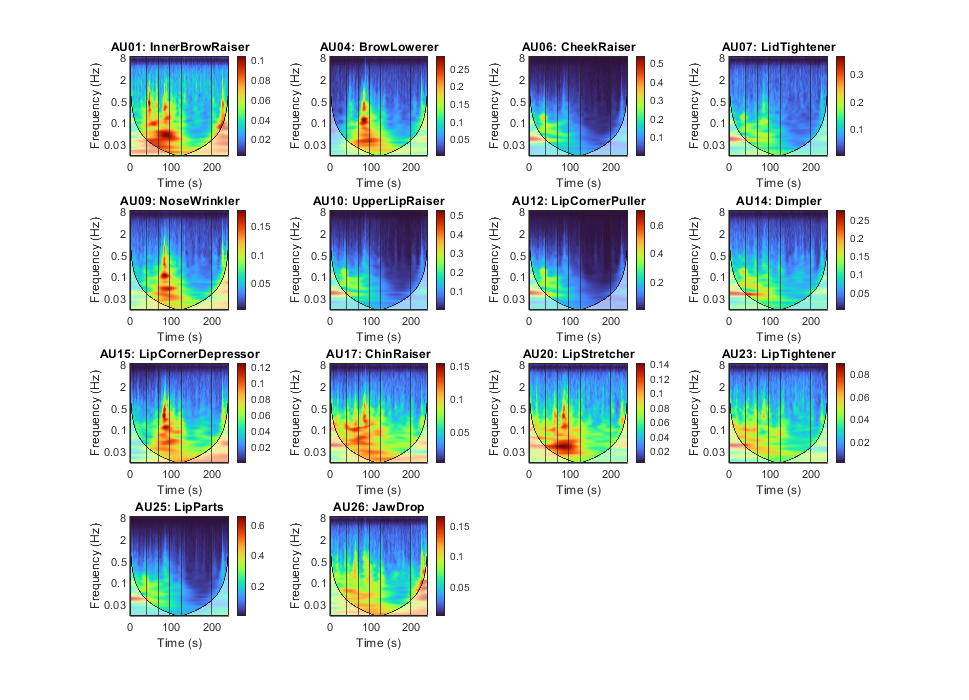

**Supplementary Figure 2.** Mean of time-frequency representation across all participants in DISFA dataset. Shading indicates the cone of influence. Vertical lines demarcate video clips described in Supplementary Table 1.

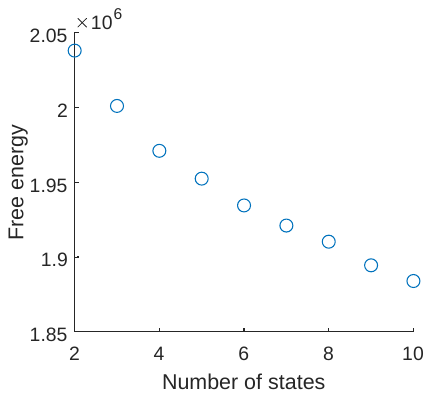

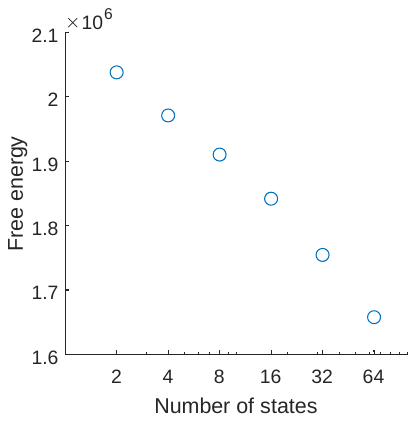

**Supplementary Figure 3.** Free energy of the Hidden Markov Model as a function of number of states. Free energy continues to decrease as the number of states increases. (Left) 2 – 10 states. (Right) 2 – 64 states, logarithmic scale.

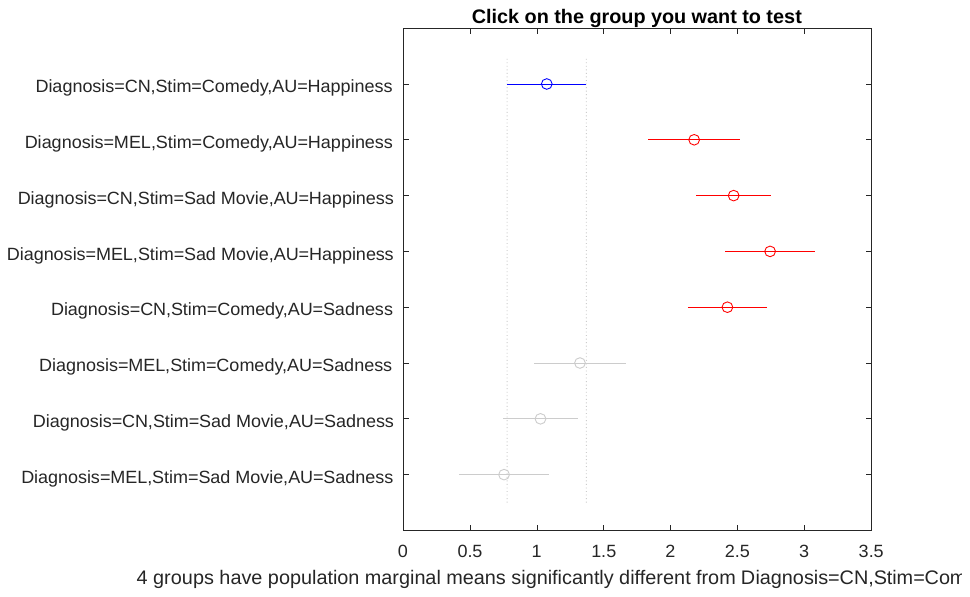

**Supplementary Figure 4.** Post-hoc comparison intervals using Tukey’s honestly significant difference criterion (p=0.05)

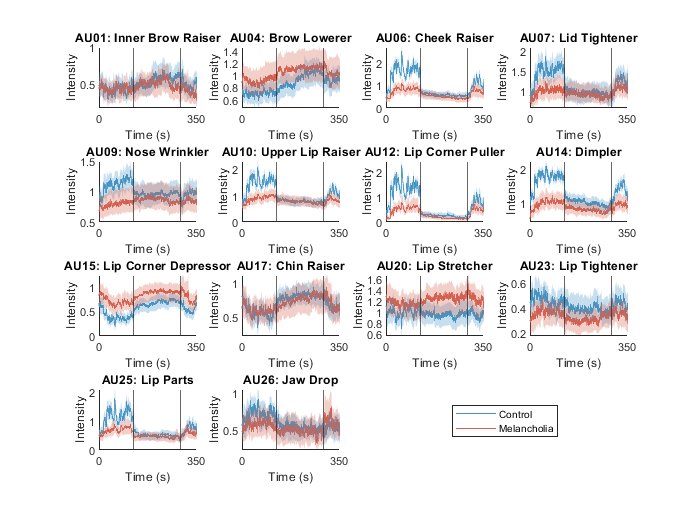

**Supplementary Figure 5.** Mean facial action unit activation in controls and melancholia for all action units. Shaded confidence bands were calculated as the 5^th^ and 95^th^ percentile in a bootstrap sample (n=100). Participants watched, in sequence, stand-up comedy, a sad video, and a funny video. Vertical lines demarcate transitions between video clips.

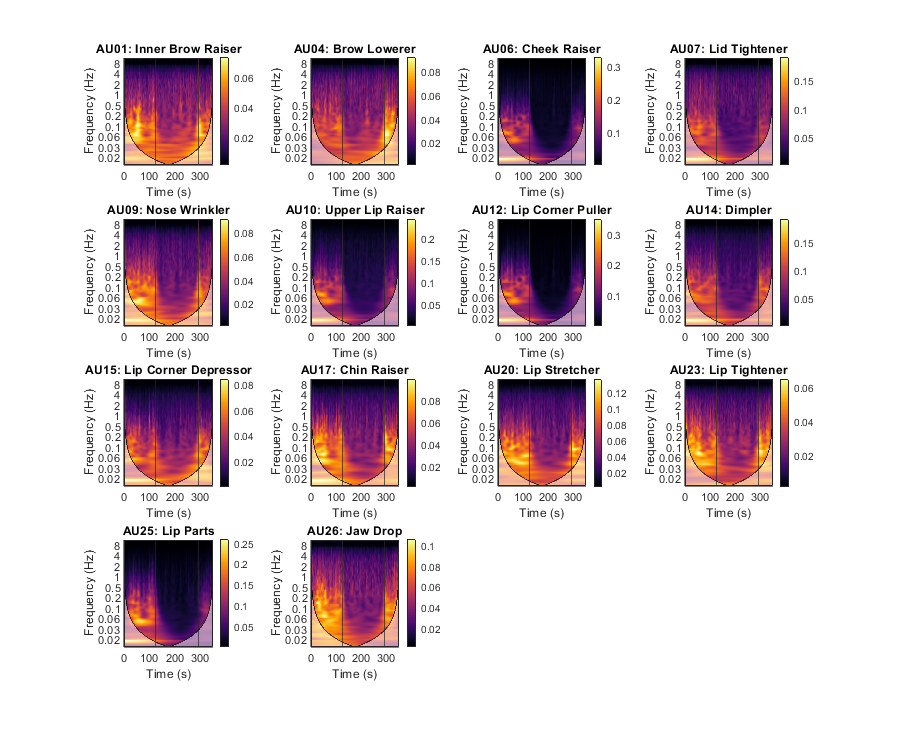

**Supplementary Figure 6.** Mean of time-frequency representation across all controls in melancholia dataset. Participants watched, in sequence, stand-up comedy, a sad video, and a funny video. Shading indicates the cone of influence. Vertical lines demarcate video clips.

| **Model** | **Number of input features** | **Individual trial accuracies** | **Mean accuracy** |
| --- | --- | --- | --- |
| Model 1: mean activation | 42 | 64% 67%, 67%, 60%, 59% | 63% |
| Model 2: Time-frequency representation in 10 frequency bands | 420 | 73%, 70%, 75%, 67%, 71% | 71% |
| Model 3A: Mean activation in 30s time chunks | 224 | 64%, 64%, 67%, 60%, 65% | 64% |
| Model 3B: Mean activation in 10s time chunks | 462 | 67%, 62%, 64%, 59%, 62% | 63% |
| Model 3C: Mean activation in 2s time chunks | 2464 | 59%, 60%, 68%, 67%, 64% | 64% |

Supplementary Table 3. Models to classify participants with melancholia from healthy controls. All models used support vector machine with Gaussian kernel, and were tested with 5-fold cross validation.

| **Parameter** | **Description** | **Value** |
| --- | --- | --- |
| K | Maximum number of HMM states | 8 |
| Order | Maximum order of multivariate auto-regressive model. 0 for no auto-regression. | 0 |
| DirichletDiag | Value of the diagonal of the prior of the transition probability matrix | 1 |
| pca | Number of top PCA components used | 10 |
| downsample | New sampling rate (Hz) | 10 |
| cyc | Maximum number of variational inference cycles | 500 |
| initcyc | Number of repetitions of the initialisation algorithm | 10 |

Supplementary Table 4. Parameters for HMM implemented in the HMM-MAR toolbox. Parameters not listed here are left to default options.

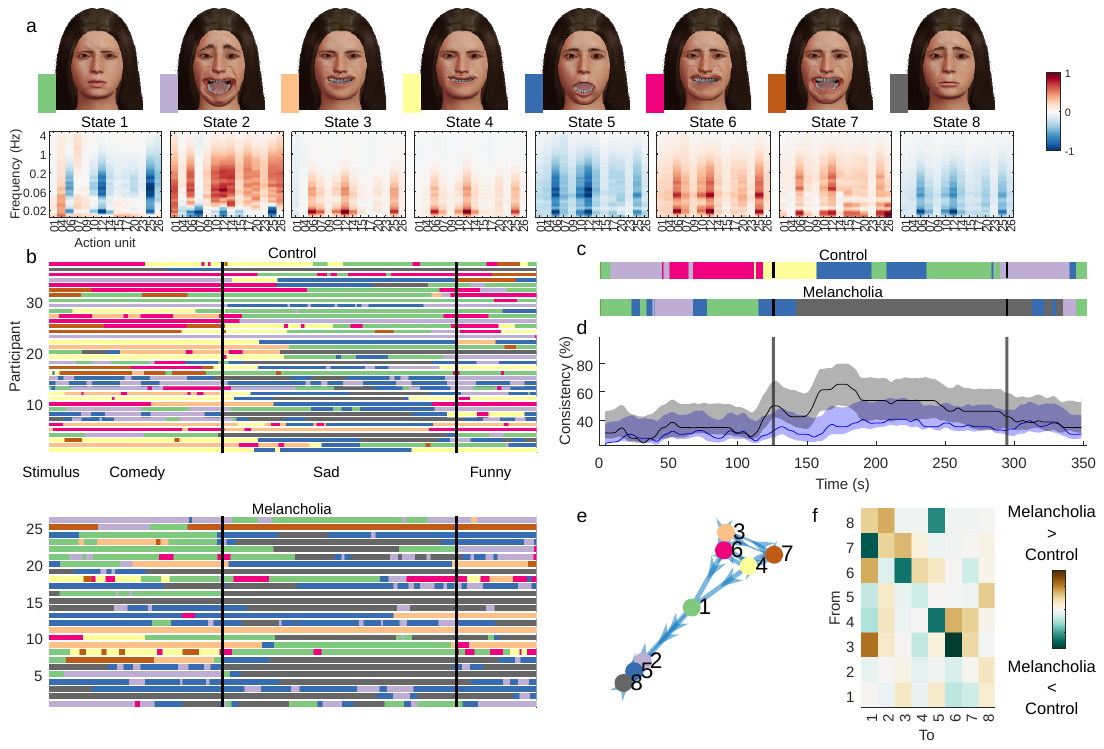

**Supplementary Figure 7.** Hidden Markov model inferred from time-frequency representation of melancholia dataset, where data are not standardised before model inference. (a) Mean of the observation model for each state. Avatar faces for each state show the relative contribution of each action unit. (b) Most likely state sequence for each participant at each time point, for controls (top) and participants with melancholia (bottom). Vertical lines demarcate stimulus clips. Without standardisation, state transitions are infrequent and transition probabilities less meaningful. (c) Most common state across participants, using a 4s sliding temporal window. (d) Proportion of participants expressing the most common state for controls (blue) and participants with melancholia (black). Shading indicates 5% and 95% bootstrap confidence bands. (e), Transition probabilities displayed as a weighted graph. Only the top 20% of transition probabilities are shown. States are positioned according to a force-directed layout where edge length is the inverse of transition probability. (f), Differences in mean transition probabilities between participants with melancholia and controls.
